# Species coexistence in resource-limited patterned ecosystems is facilitated by the interplay of spatial self-organisation and intraspecific competition

**DOI:** 10.1101/2020.01.13.903179

**Authors:** Lukas Eigentler

**Affiliations:** Division of Molecular Microbiology, School of Life Sciences, University of Dundee, Dundee DD1 5EH, United Kingdom; Division of Mathematics, School of Science and Engineering, University of Dundee, Dundee DD1 4HN, United Kingdom; Maxwell Institute for Mathematical Sciences, Department of Mathematics, Heriot-Watt University, Edinburgh EH14 4AS, United Kingdom

**Keywords:** pattern formation, competition and coexistence, banded vegetation, periodic travelling waves, scale-dependent feedback, numerical continuation

## Abstract

The exploration of mechanisms that enable species coexistence under competition for a sole limiting resource is widespread across ecology. Two examples of such facilitative processes are intraspecific competition and spatial self-organisation. These processes determine the outcome of competitive dynamics in many resource-limited patterned ecosystems, classical examples of which include dryland vegetation patterns, intertidal mussel beds and Sub-alpine ribbon forests. Previous theoretical investigations have explained coexistence within patterned ecosystems by making strong assumptions on the differences between species (e.g. contrasting dispersal behaviours or different functional responses to resource availability). In this paper, I show that the interplay between the detrimental effects of intraspecific competition and the facilitative nature of self-organisation forms a coexistence mechanism that does not rely on species-specific assumptions and captures coexistence across a wide range of the environmental stress gradient. I use a theoretical model that captures the interactions of two generic consumer species with an explicitly modelled resource to show that coexistence relies on a balance between species’ colonisation abilities and their local competitiveness, provided intraspecific competition is sufficiently strong. Crucially, the requirements on species’ self-limitation for coexistence to occur differ on opposite ends of the resource input spectrum. For low resource levels, coexistence is facilitated by strong intraspecific dynamics of the species superior in its colonisation abilities, but for larger volumes of resource input, strong intraspecific competition of the locally superior species enables coexistence. Results presented in this paper also highlight the importance of hysteresis in understanding tipping points, in particular extinction events. Finally, the theoretical framework provides insights into spatial species distributions within single patches, supporting verbal hypotheses on co-existence of herbaceous and woody species in dryland vegetation patterns and suggesting potential empirical tests in the context of other patterned ecosystems.

## 1 Introduction

A classical result of coexistence theory states that species coexistence in ecological and biological systems cannot occur if species compete for a single limiting resource (Hutchinson 1959; Levin 1970; R. MacArthur and Levins 1964; Tilman 1982); this is a consequence of the competitive exclusion principle (Gause 1934). Indeed, according to Tilman’s *R*^∗^ rule, the species that is able to depress the limiting resource to its lowest level is expected to exclude all competitors (Tilman 1982). However, species coexistence occurs in many ecosystems despite competition for a single resource (e.g. (Babarro and Comeau 2014; Porri et al. 2016; Reise et al. 2017; Seghieri et al. 1997; Valentin et al. 1999)). Hence, this suggests that other processes prevent resource monopolisation by one species and thus enable coexistence in these ecosystems (McPeek 2012). One such mechanism that has been singled out as a common facilitator of species coexistence is intraspecific competition (e.g. (Chesson 2000; R. MacArthur 1970; McPeek 2012; Tilman 1982)). The inclusion of self-limitation due to negative density-dependent effects is an integral part of investigating coexistence through Lotka-Volterra competition models (see (Chesson 2000) for a discussion). More recently, such dynamics have also been added to Rosenzweig-MacArthur-type models in which interactions of consumer species with a sole limiting resource are explicitly modelled (McPeek 2012). The impact of self-limitation on consumer competition dynamics is that, if sufficiently strong, it limits the abundance of each species so that no single species is able to monopolise the resource in the sense of Tilman’s *R*^∗^ rule. This allows the limiting resource to be shared among two or more species and thus enables coexistence (McPeek 2012).

Another classical theory of competition and coexistence theory is the stabilisation of species coexistence in resource limited environments due to a trade-off between species’ dispersal and competitive abilities (Horn and R. H. MacArthur 1972; Levins and Culver 1971). Species that are outcompeted locally by a locally superior competitor can persist through rapid invasion of uncolonised areas (Tilman 1994). Thus, such a balance is attributed to enable coexistence on a regional or metacommunity scale through spatial segregation in a patchy but connected environment, despite competitive exclusion taking place on a local, single-patch scale (Gravel et al. 2010; Hassell et al. 1994; Horn and R. H. MacArthur 1972; Levins and Culver 1971; Tilman 1994).

More recently, theory on spatial self-organisation has utilised classical results based on the colonisation-competition trade-off to explain species coexistence on a local scale in an otherwise homogeneous environment (Cornacchia et al. 2018; Eigentler and Sherratt 2020c). As opposed to local intraspecific competition, self-organisation principles are characterised by positive density-dependent effects on short spatial scales that occur in combination with long-range competition for resources (Rietkerk and van de Koppel 2008). The formation of spatial patterns of consumer species due to such scale-dependent feedbacks can create spatial heterogeneities in environmental conditions (e.g. resource availability) in otherwise homogeneous environments (Cornacchia et al. 2018; Eigentler and Sherratt 2020c). Such heterogeneous landscapes lead to spatial separation of competitive and facilitative interactions between different consumer species, which creates a balance that enables species coexistence without the need for spatial segregation (Cornacchia et al. 2018). This mechanism is not unique to self-organised consumer communities, but has also been attributed to enrich species richness in other ecological systems where the underlying spatial heterogeneities are not induced by the species themselves (van de Koppel et al. 2006).

Notably, in isolation, both intraspecific competition dynamics and spatial self-organisation principles have a stabilising impact on species coexistence, despite their fundamental differences in terms of population growth. Intraspecific competition is typically associated with negative density dependence, while spatial self-organisation is characterised by positive density dependence. In this paper I aim to assess the impact of the interplay between intraspecific competition and spatial self-organisation principles on species coexistence in resource limited ecosystems by unifying them in a theoretical framework. Particular focus is put on their role under severe environmental stress, as spatial self-organisation principles are known to have a strong impact on populations under such conditions (Rietkerk and van de Koppel 2008).

Indeed, spatial self-organisation principles become particularly striking and visible to the naked eye if environmental conditions are sufficiently harsh to necessitate a population’s separation into colonised and uncolonised areas (Rietkerk and van de Koppel 2008). Examples include but are not limited to dryland vegetation patterns (Deblauwe et al. 2012; Gandhi et al. 2019; Seghieri et al. 1997; Valentin et al. 1999), mussel beds in intertidal regions (Gascoigne et al. 2005; Liu et al. 2014; van de Koppel et al. 2005) and ribbon forests along tree lines in Subalpine ecosystems (Bekker 2005; Bekker and Malanson 2008; Hiemstra et al. 2002; Hiemstra et al. 2006). Despite occurring in distant geographical locations and under fundamentally different environmental conditions, these ecosystems are united by their self-organisation into alternating patches of colonised and uncolonised areas to combat severe environmental stress. In the context of dryland vegetation patterns, this is underpinned by a positive feedback between local vegetation growth and water redistribution towards areas of high biomass allowing a spatially structured population to persist under arid conditions (Gandhi et al. 2019; Rietkerk and van de Koppel 2008). Similarly, spatial aggregation of trees along tree lines in form of stripes perpendicular to prevailing wind conditions enhances seedling establishment due to protection from wind-driven snow accumulation (Bekker 2005; Bekker and Malanson 2008; Hiemstra et al. 2002; Hiemstra et al. 2006). In mussel beds, facilitation in densely populated regions occurs due to a reduction in wave disturbance and protection from predators (Bertness and Grosholz 1985; Côté and Jelnikar 1999; Hunt and Scheibling 2001). Combined with scarcity of the mussels’ main food source, algae (Dolmer 2000), this creates a pattern-inducing feedback. Moreover, despite the environmental stress associated with these ecosystems, occurrence of species co-existence on the scales of single colonised patches have been reported. In dryland vegetation patterns, herbaceous species typically coexist with shrubs or trees (Seghieri et al. 1997; Valentin et al. 1999); ribbon forests in the Rocky Mountains feature spruces (*Picea engelmannii*) and firs (*Abies lasiocarpa*) (Bekker and Malanson 2008; Hiemstra et al. 2006); and Pacific oysters (*Magallana gigas*) and blue mussels (*Mytilus edulis*) form mussel beds in the Wadden Sea (Reise et al. 2017), with coexistence of different species being reported from other geographical locations (Babarro and Comeau 2014; Porri et al. 2016).

Many patterned ecosystems have been subject to theoretical models to complement field and experimental studies, including mussel beds (Bennett and Sherratt 2018a; Holzer and Popović 2017; Liu et al. 2012; Liu et al. 2014; Shen and Wei 2020; Sherratt and Mackenzie 2016; van de Koppel et al. 2005) and ribbon forests (Hiemstra et al. 2002; Hiemstra et al. 2006). The quantity of theoretical research on these particular ecosystems is dwarfed by the vast number of mathematical models on dryland vegetation patterns (see (Borgogno et al. 2009; Gandhi et al. 2019; Martinez-Garcia and Lopez 2018; Zelnik et al. 2013) for reviews). These approaches include models to investigate species coexistence in dryland vegetation patterns which, to this date, have been unable to capture species coexistence as a stable model outcome in the absence of species-specific assumptions, such as a distinction between species inducing pattern formation and species unable to form a spatial pattern (Baudena and Rietkerk 2013; Nathan et al. 2013), or species’ adaptation to different soil moisture niches (Callegaro and Ursino 2018; Ursino and Callegaro 2016). Models attempting to overcome this shortfall by considering species that do not fundamentally differ in their interaction with the environment only capture species coexistence as a transient feature (Eigentler and Sherratt 2019; Gilad et al. 2007) or under less severe environmental stress (Eigentler and Sherratt 2020c).

Indeed, it has been shown that self-organisation is a sufficient mechanism to enable coexistence in continuously colonised states, where resource availability is sufficiently high to prevent segregation into colonised and bare patches (Eigentler and Sherratt 2020c). Crucially however, it is also argued that other processes must be involved in the facilitation of coexistence in patterned ecosystems under severe environmental stress if species do not differ in their functional responses to the environment. This hypothesis is based on a description of the resource-consumer dynamics of two species (or two groups of species) whose difference from each other only manifests itself quantitatively through different parameter estimates of basic properties (e.g. growth and death rates) but not qualitatively through different functional responses to the environment. In this setting, coexistence occurs if one (group of) species is superior in its colonisation abilities but is outcompeted locally by a second (group of) species. Consequently, the former are referred to as *colonisers* and the later as the *locally superior species*, a terminology that will be carried over to this paper. The argument is related to theory on coexistence induced by a trade-off between competition and dispersal (e.g. (Horn and R. H. MacArthur 1972; Levins and Culver 1971)), with the crucial difference that spatial self-organisation enables coexistence without the need for spatial segregation between the coexisting species. This mechanism only gives insights into the coexistence of both species groups. Species richness within those groups may occur due to other facilitative processes that are independent of spatial self-organisation principles (see (Godwin et al. 2020) for a review on both historic and more recent advances). For brevity, I refer to such species groups as single species in the remainder of the paper.

This previous study functions as the baseline case of the unification of spatial self-organisation and intraspecific competition dynamics in coexistence theory of patterned ecosystems presented in this paper. Even though the underlying modelling framework’s origin is the plant-water population dynamics in drylands, I argue that its description of key ecosystem processes and feedbacks is sufficiently general to apply to a whole host of other patterned ecosystems, including but not being limited to those of mussel beds and ribbon forests. I show that the inclusion of self-limitation is crucial to capture coexistence of a coloniser species with a locally superior species under severe environmental stress. Moreover, the importance of spatial self-organisation in the coexistence of consumer species in resource limited ecosystems is reinforced by the result that strong intraspecific competition of different species facilitates coexistence at different extrema of the resource input spectrum. Finally, this unified approach also sheds light on species distribution within single patches, thereby confirming verbal arguments that, in the context of vegetation patterns, the stripes’ uphill migration on sloped terrain is indeed driven by the coloniser species (Seghieri et al. 1997).

This paper focusses on the ecological implications obtained through a comprehensive model analysis of the system presented in Sec. 2 below. As such, only stable model outcomes are considered. Nevertheless, the mathematical model also admits unstable solutions, an analysis of which aids greatly to the understanding of the bifurcations in the system. A comprehensive bifurcation analysis aimed at a more mathematics-oriented readership and an analysis of the impact of the interplay of intraspecific competition dynamics and spatial self-organisation in single-species ecosystems are presented in a separate publication (Eigentler 2020) that puts particular emphasis on dryland vegetation patterns.

## 2 Model & Methods

The mathematical model used to investigate the impact of the interplay of intraspecific competition and spatial self-organisation on species coexistence in patterned ecosystems is based on the Klausmeier model (Klausmeier 1999), one of the prototype toy models used in theoretical studies of dryland vegetation patterns (e.g. (Bastiaansen et al. 2018; Carter and Doelman 2018; Sherratt 2010; Sherratt 2005; Sherratt 2011; Sherratt 2013b; Sherratt 2013c; Sherratt 2013d; Siteur et al. 2014)). Despite its origin, I argue that its deliberately simple description of the self-organisation principle as well as other processes extends its applicability to a generic setting, as I outline in Section 2.1 below. Moreover, it provides a rich framework for model extensions (e.g. (Bennett and Sherratt 2018b; Consolo and Valenti 2019; Eigentler and Sherratt 2020a; Eigentler and Sherratt 2018; Eigentler and Sherratt 2020b; Fernandez-Oto et al. 2019; Gandhi et al. 2018; Marasco et al. 2014; Siero 2018; Siero et al. 2019; Wang and Zhang 2018; Wang and Zhang 2019)). One recent extension has introduced a second consumer species to the system, based on the assumption that both species only differ from each other quantitatively in their basic parameters but not qualitatively in any of their functional responses (Eigentler and Sherratt 2019; Eigentler and Sherratt 2020c). This model forms the basis of the theoretical framework presented below.

### 2.1 Model details

The focus of this paper lies on the characterisation of the impact of the interplay of intraspecific competition and spatial self-organisation dynamics on species coexistence. The modelling framework presented in (Eigentler and Sherratt 2019; Eigentler and Sherratt 2020c) only accounts for the facilitative impact of the latter and assumes that the per capita growth rates of consumer species are given by

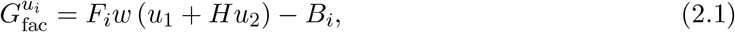

where *u*_1_ = *u*_1_(*x, t*) and *u*_2_ = *u*_2_(*x, t*) denote the nondimensional (see the supplementary material for a nondimensonalisation) consumer densities at time *t* ≥ 0 at the space point *x* ∈ ℝ. The per capita growth rate not only increases with the resource density *w* = *w*(*x, t*), but also with both consumer densities *u*_1_ and *u*_2_ (Fig. 2.1). This positive density-dependence represents the short-range facilitative effects of consumers on each other which is caused, for example, by increases of soil permeability in vegetated areas in drylands or by enhancements in tree seedling establishment in ribbon forests due to protection from snow by adult trees. The facilitative impact may differ between the two species and is accounted for by the nondimensional constant *H*. The nondimensional constants *F*_*i*_ describe the species’ resource-to-biomass conversion capabilities. Note that the per capita growth rate may be negative, as it includes a baseline mortality, assumed to occur at constant rates *B*_*i*_. However, this does not necessitate a clear distinction between birth and death in the two terms in (2.1), as the facilitation term may also represent reductions in mortality instead of enhancements of growth, for example increased protection from wave dislodgement or predation in dense mussel communities.

**Figure 2.1:**
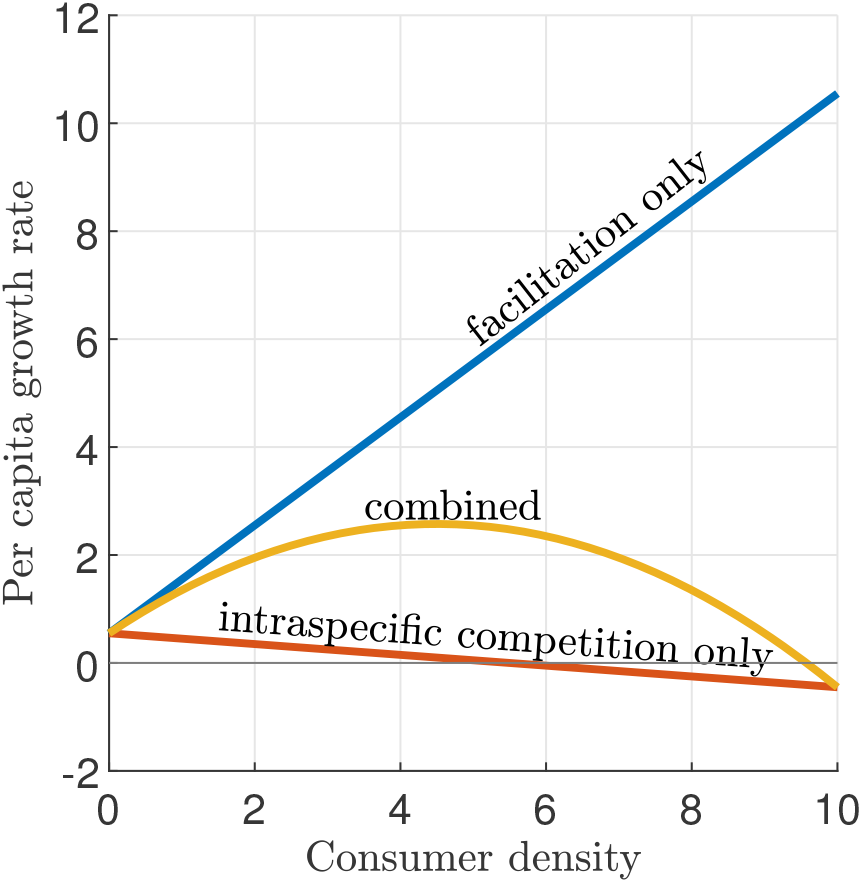
Per capita consumer growth rates. Sketches of a consumer species’ per capita growth rate are shown for a growth regime featuring facilitation only (blue), intraspecific competition only (red) and an interplay of facilitation and intraspecific competition (yellow). Facilitation is associated with increases in per capita growth rates for increasing consumer density, while intraspecific competition results in decreases of the per capita growth rate. The interplay between intraspecific competition and facilitation results in dominance of facilitation for low consumer densities and inhibition due to intraspecific competition at high consumer densities. Note that due to the inclusion of the baseline mortality rate *B*_*i*_ in the per capita growth rate of consumer species *u*_*i*_, the growth rate becomes negative at a consumer density lower than the species’ carrying capacity in the absence of mortality (*k*_*i*_). For this visualisation, resource abundance, density a competitor species and the consumer’s base mortality are assumed to be constant.

The per capita growth rate (2.1) neglects any intraspecific competition dynamics and thus leads to unrealistically high growth rates at high consumer densities (Fig. 2.1), which may result in biologically unrealistic model solutions (Bennett and Sherratt 2018b). To include self-limitation by consumer species, I adjust the per capita growth rates (2.1) to

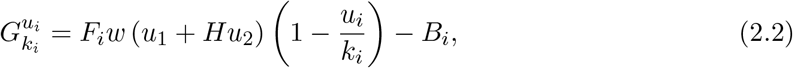

where the parameter *k*_*i*_ denotes the carrying capacity of species *u*_*i*_ in the absence of the baseline mortality *B*_*i*_. Notably, due to the nonzero baseline mortality *B*_*i*_, the actual carrying capacity of the consumer species *u*_*i*_ is less than or equal *k*_*i*_ (Fig. 2.1). Thus, *k*_*i*_ is referred to as the reciprocal strength of the self-limitation among consumer species *u*_*i*_, rather than its carrying capacity. The inclusion of the inhibitory 1−*u*_*i*_*/k*_*i*_ term in the per capita growth rate is motivated by its frequent usage in the modelling of population dynamics and can be dated back to the 19th century (Bacaër 2011; Verhulst 1838). In the absence of facilitative self-organisation dynamics and the baseline mortality, it is typically used to describe logistic growth through a linear decrease of the per capita growth rate with increasing consumer density (Fig. 2.1). It is noteworthy that in this context, self-limitation does not refer to intraspecific competition for the limiting resource, as these dynamics are explicitly accounted for in the system. Instead, the strength of intraspecific competition depends on other factors, which limit the total biomass a species can reach in a fixed area, such as the maximum biomass of a single individual (Nathan et al. 2013), or, in the context of plant ecosystems, the decomposition of plant litter leading to an accumulation of autotoxic soil compounds (Bonanomi et al. 2011; Mazzoleni et al. 2007). By contrast, interspecific competition is only assumed to occur through depletion of the limiting resource. The inhibitory 1 − *u*_*i*_*/k*_*i*_ term in the per capita growth rate is thus independent of the competitor species *u*_*j*_, This provides a stark contrast to other commonly used competition models (e.g. (Chesson 2000; Eppinga et al. 2018)), in which such a term typically represents competition for resources (e.g. space) that are not explicitly accounted for in the model. As competition for limiting resources is typically singled out as a major obstacle to species coexistence, the model presented below neglects any additional interspecific competition dynamics that could lead to other coexistence mechanisms and thus solely focusses on the potential for species coexistence to occur despite competition for a sole limiting resource.

Combined, the interplay of spatial self-organisation and intraspecific competition results in a nonlinear relation between per capita growth rate and consumer density that is underpinned by facilitation while the consumer is scarce and inhibition as the consumer becomes more abundant in the ecosystem (Fig. 2.1).

Replacing the plant growth rates (2.1) in the multispecies model presented in (Eigentler and Sherratt 2019; Eigentler and Sherratt 2020c) by (2.2) yields the theoretical framework used to characterise the impact of the interplay of spatial self-organisation and intraspecific competition on species coexistence. Suitably nondimensionalised (see supplementary material), it is

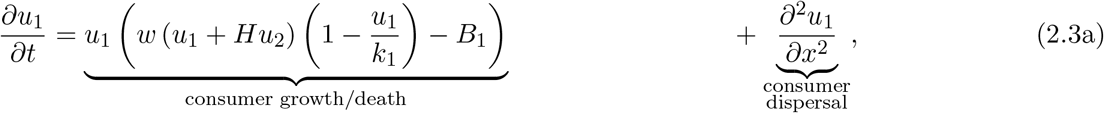

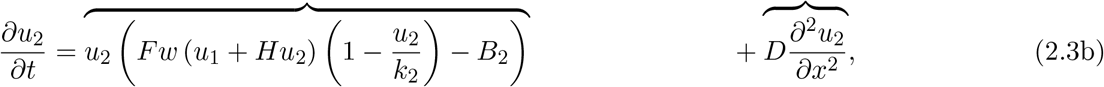

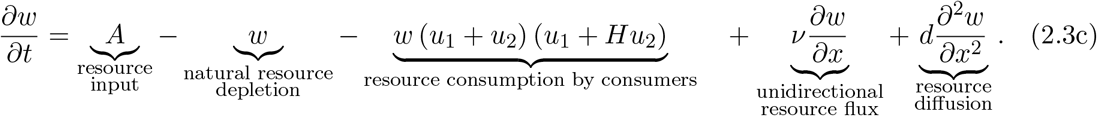

The density *w* is referred to as the limiting resource (e.g. water in the context of dryland vegetation, algae in mussel beds) throughout the paper, but, in more general terms, should be interpreted as a proxy for environmental conditions (e.g. a reciprocal measure of snow abundance in ribbon forests).

The resource is added to the system at a constant rate *A* (rainfall in dryland vegetation, algae influx from upper water layers in mussel beds, melting of snow in ribbon forests) and is removed at a constant rate due to processes independent of the consumer species (water evaporation and drainage, natural algae death, snowfall). Resource consumption by both consumer species is represented by the third term on the right hand side of (2.3c). The rate of resource consumption does not only depend on the total consumer density (*u*_1_ + *u*_2_), but also on the enhancement of growing conditions due to facilitation in areas of high biomass (*u*_1_ + *Hu*_2_), as discussed above (enhancement of water infiltration into the soil, reduction of mussel mortality, enhanced tree seedling survival). Resource consumption remains unaffected by the additional intraspecific competition dynamics that cause a reduction in consumer growth as the consumer density increases. While this logistic-type term reduces the consumers’ resource-to-new-biomass conversion efficiency, it accounts for processes that are independent of the resource competition dynamics and thus has no impact on the rate of resource consumption by existing consumer biomass. Finally, all three densities diffuse and the resource is assumed to undergo additional unidirectional flux, described by an advection term in (2.3c). This assumption holds true in many ecosystems; for example, it accounts for water flow downhill on gentle slopes on which dryland vegetation patterns can be found (Valentin et al. 1999); represents algae movement due to tidal flows (van de Koppel et al. 2005); or snow displacement along the prevailing wind direction in Subalpine ribbon forests (Bekker 2005). This form of resource flux is commonly accredited with the appearance of spatial patterns in the form of regular stripes perpendicular to the direction of the flux (Hiemstra et al. 2002; Valentin et al. 1999; van de Koppel et al. 2005).

It is possible to capture such regular banded patterns by using a one-dimensional spatial domain instead of an ecologically more intuitive two-dimensional domain in the modelling framework. This is common practice in the study of stripe patterns (e.g. (Gandhi et al. 2020; Klausmeier 1999; Sherratt 2016)) as outcomes are similar to those obtained on a two-dimensional spatial domain. However, it is important to emphasise that this may lead to an overestimation of a pattern’s resilience to increases in environmental stress, as it cannot capture the break-up of banded patterns into spot patterns (Sewalt and Doelman 2017; Siero et al. 2015). Solution profiles of (2.3) thus represent transversal cuts along the direction of the resource flux. Nevertheless, I argue that results presented in this paper can be extended to ecosystems that lack such a unidirectional resource flux (e.g. dryland vegetation patterns on flat terrain), as is discussed in Section 4. The nondimensional diffusion parameters *D* for species *u*_2_ and *d* for water, as well as the advection speed *ν* compare the respective dimensional parameters to the diffusion coefficient of species *u*_1_.

In the following, I assume that *u*_1_ and *u*_2_ represent a *coloniser species* and a *locally superior species*, respectively. The distinction between a coloniser and a locally superior species yields qualitative assumptions on the model parameters (Eigentler and Sherratt 2020c). The coloniser both grows and dies at a faster rate (*F <* 1, *B*_1_ *> B*_2_, but see comment below), has a stronger impact on the spatial self-organisation principle (*H <* 1) and disperses faster (*D <* 1). Details on how such qualitative relations can be obtained in the context of dryland vegetation despite the lack of empirical data can be found in (Accatino et al. 2010; Eigentler and Sherratt 2020c; Klausmeier 1999; Mauchamp et al. 1994), but I emphasise that results presented in this paper are robust to changes in parameter values that do not violate the qualitative assumptions. The locally superior species is assumed to outcompete the coloniser species in a spatially uniform setting in the absence of any self-limitation. This requires *B*_2_ *< FB*_1_ (Eigentler and Sherratt 2019) and I assume that this relation holds throughout the paper.

### 2.2 Model analysis

Typically (but see Fig. 3.2 (d) and Fig. S5.3 in the supplementary material for one exception), both single-species solutions and coexistence solutions of (2.3) either occur in a spatially constant configuration or as regular spatial patterns. In the latter case, they are periodic travelling waves, i.e. spatially periodic functions that move in the direction opposite to the unidirectional resource flux at a constant speed. Thus, they are periodic in both space and time and are therefore referred to as *spatiotemporal patterns*. The migration speed is an emergent property of model solutions, but can be made explicit through a change of coordinates (see supplementary material). This facilitates a bifurcation analysis (the study of qualitative changes to the solution structure under variations of system parameters), which is performed by a combination of analytical tools and numerical continuation (see supplementary material). In particular, the stability calculations of the periodic travelling wave solutions of the system are performed using a numerical continuation method for the calculation of essential spectra by Rademacher et al. 2007.

Numerical simulations of the PDE system (2.3) are obtained as follows. The spatial domain is discretised into a number of equidistant points (here 2^9^ points) and spatial derivatives are approximated by finite differences (forward and central finite differences for first and second derivatives, respectively) on the discretised spaital domain. This results in a system of coupled ordinary differential equations (ODEs), which is solved by Matlab’s ODE solver ode15s. While both the analytical results and the numerical continuation algorithms are based on the assumption that the spatial domain is of an infinite size, numerical simulations of the system require a finite spatial domain. Boundary conditions are chosen to be periodic as they provide the best approximation to an infinite domain length. The length of the spatial domain imposes a restriction on the possible wavelengths of a spatial pattern, as the domain length is required to be a multiple of the pattern’s wavelength. Except this restriction, the length of the spatial domain bears no significant influence on the model dynamics. As numerical simulations of the model are only used for visualisation purposes but not for means of model analysis throughout the paper, the length of the spatial domain varies between different realisations, depending on the intended purpose of the visualisation. Model simulations are initialised from solutions obtained through numerical continuation.

The spatial distribution of species in a single consumer band is quantified by the linear correlation between both species’ solution components. The linear correlation is given by

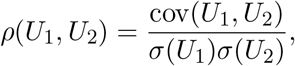

where *U*_1_ and *U*_2_ are two vectors obtained by discretising the spatial domain in space and evaluating the consumer densities *u*_1_ and *u*_2_ on this mesh. Here, cov(*·, ·*) denotes the covariance of two vectors, and *σ*(*·*) the standard deviation. The linear correlation satisfies −1 ≤ *ρ*(*U*_1_, *U*_2_) ≤ 1, and a larger correlation corresponds to a more in-phase-like appearance of both consumer patterns. Numerical continuation of model solutions using AUTO-07p (Doedel et al. 2012) allows for an exhaustive calculation of the linear correlation in the parameter space.

## 3 Results

### 3.1 Spatially uniform coexistence

Depending on the strength of resource input *A*, the multispecies model (2.3) has up to four biologically relevant spatially uniform equilibria: an extinction steady state that is stable in the whole parameter space; a single-species equilibrium for each consumer species; and a coexistence state. Under sufficiently high resource input levels, the single-species equilibria are stable in the absence of a second species, but may become unstable if a competitor is introduced. Stability to the introduction of a second species requires both sufficiently weak intraspecific competition and a local average fitness higher than that of the competitor. Moreover, the stability regions of both single-species equilibria do not overlap and it is straightforward to determine the species of higher local average fitness based on their parameter values. In particular, only changes to growth and mortality rates, but not variations in the strength of the intraspecific competition can change which species is of higher local average fitness (see supplementary material for more details).

As intraspecific competition of the locally superior species increases, its single-species equilibrium loses stability to the coexistence equilibrium (Fig. 3.1). In other words, spatially uniform coexistence occurs if intraspecific competition among the locally superior species is sufficiently strong compared to the interspecific competition for the limiting resource. Strong intraspecific competition reduces equilibrium densities and hence renders the locally superior species unable to utilise all of the available resource. This allows for the invasion of a second species and thus facilitates coexistence. The single species equilibrium of the locally superior species loses its stability as it intersects with the coexistence equilibrium (see supplementary material). Thus, the transition from a single-species state to coexistence is a gradual process, characterised by a gradual increase of the coloniser’s density from zero as the intraspecific competition strength of the locally superior species passes through the critical transition threshold. Moreover, if intraspecific competition among the locally superior species is sufficiently strong, then its single-species spatially uniform equilibrium is always unstable, either to the introduction of a competitor species or to the extinction steady state. Thus, under such circumstances, increases in resource stress (i.e. increases in the strength of interspecific competition) lead to a direct transition from coexistence to extinction in the spatially uniform setting (Fig. 3.1). Strong intraspecific competition also enables the locally superior species to persist in coexistence with a second species at resource input levels that are unsustainable in the absence of a increasing intraspecific competition competitor species (Fig. 3.1). This facilitative effect occurs if the competitor species is able to maintain a spatially uniform state at a lower resource input, which also has a favourable impact on the minimum resource requirements of the coexistence equilibrium. More information on the derivation of these stability boundaries are presented in the supplementary material.

**Figure 3.1:**
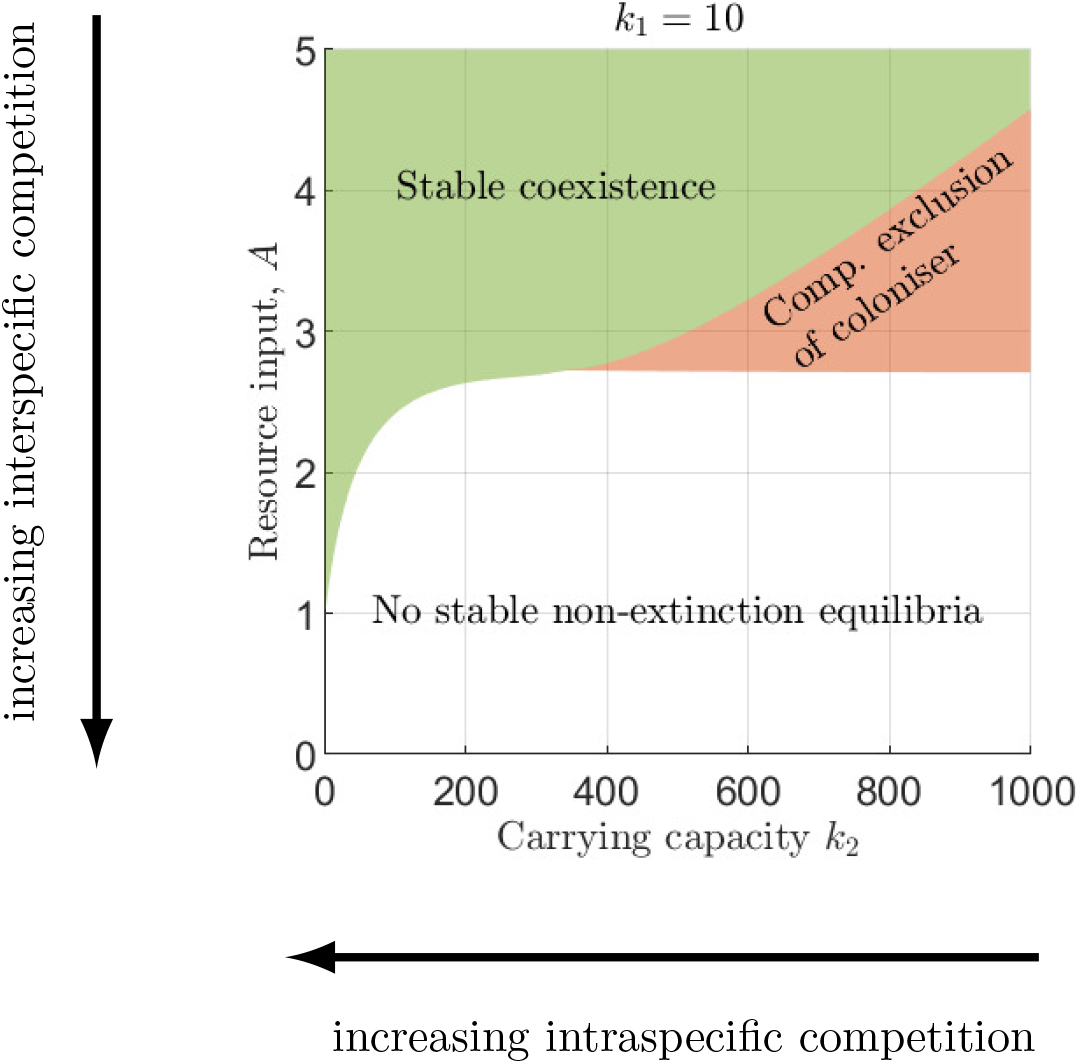
Stability of spatially uniform equilibria in the nonspatial model. Stability regions of the coexistence equilibrium and the equilibrium of the locally superior species are shown under changes to interspecific (resource input parameter *A*) and intraspecific (carrying capacity *k*_2_) competition. The equilibrium of the coloniser species is always unstable and the extinction steady state is stable in the whole parameter region (not shown). Parameter values are *B*_1_ = 0.45, *B*_2_ = 0.049, *F* = 0.11, *H* = 0.11. For a detailed overview on the stability boundaries underpinning this visualisation, see Fig. S3.1 in the supplementary material.

**Figure 3.2:**
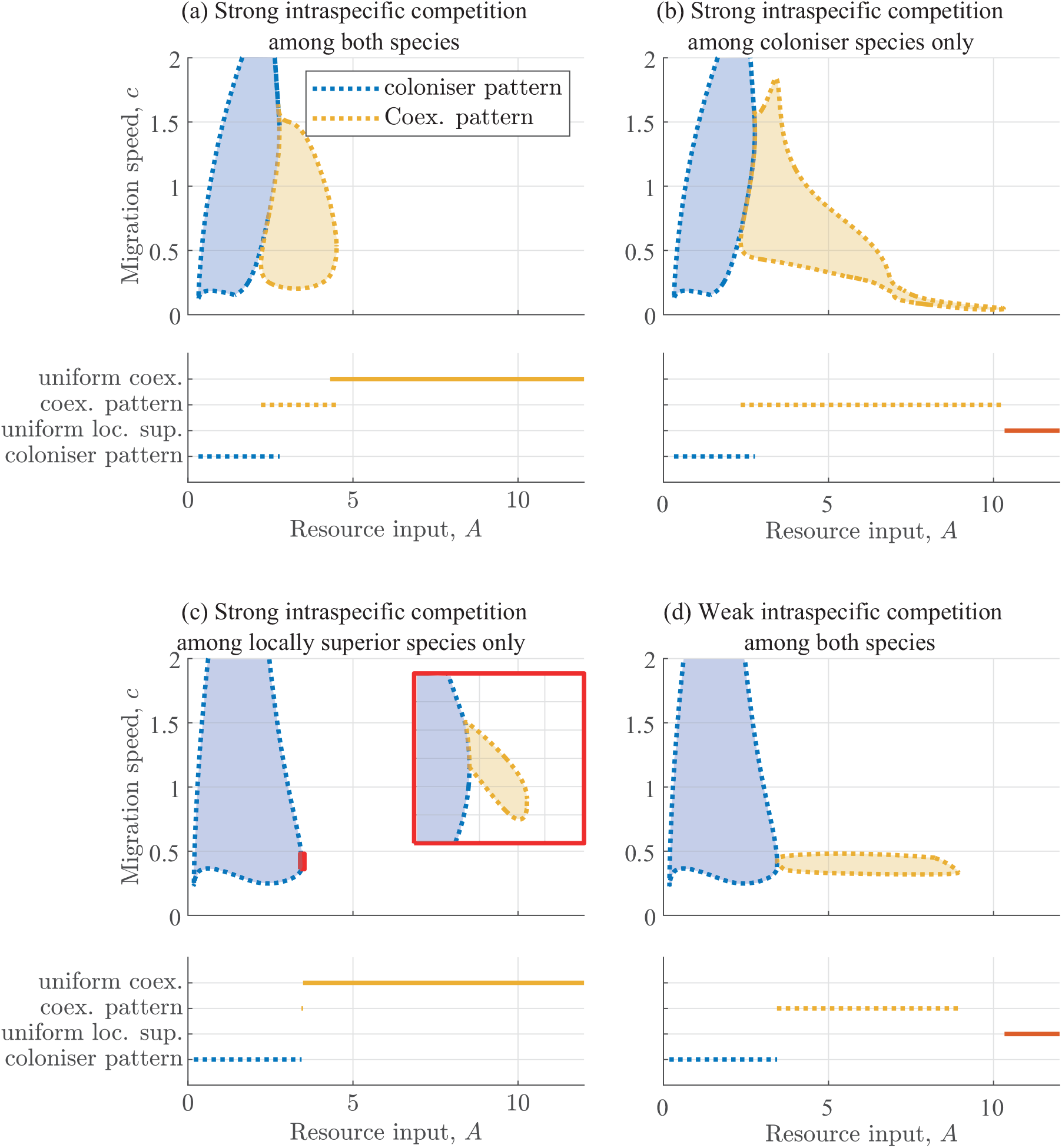
Overview of stable model outcomes for different combinations of species’ intraspecific competition strengths. The lower panel in each quadrant provides a visualisation of the resource input ranges in which spatially uniform (solid lines) and spatially patterned solutions (dotted lines) are stable model solutions. The shaded areas in the top panel shows the stability regions of the patterned states in the (*A, c*) parameter plane and thus also provides information on the uphill migration speed of stable patterns. Increases in intraspecific competition strength of the locally superior species stabilises spatially uniform coexistence for lower precipitation volumes, while strong intraspecific competition among the coloniser species promotes patterned coexistence. The intraspecific competition strengths are *k*_1_ = *k*_2_ = 10 in (a), *k*_1_ = 10, *k*_2_ = 10^4^ in (b), *k*_1_ = 10^4^, *k*_2_ = 10 in (c) and *k*_1_ = *k*_2_ = 10^4^ in (d), while other parameter values are *B*_1_ = 0.45, *B*_2_ = 0.049, *F* = *H* = *D* = 0.11, *ν* = 182.5 and *d* = 500 in (a)-(d).

Interspecific competition through depletion of the limiting resource is quantified by the resource input parameter *A* (Fig. 3.1 and 3.2). The characterisation of the strength of interspecific competition through such a proxy is necessitated by the explicit modelling of the resource dynamics. This makes it impossible to directly quantify the competitive impact of one species on the other. In particular, this does not allow for a quantitative comparison with the strength of intraspecific competition dynamics, quantified by the carrying capacities *k*_1_ and *k*_2_, as is possible in Lotka-Volterra-type models (e.g. (Chesson 2000)). The strength of intraspecific competition among the locally inferior species has no significant impact on the occurrence of spatially uniform coexistence.

For each of the non-extinction equilibria of (2.3), there exists a complementary unstable steady state with the same species composition. Their instability for all biologically realistic parameter values causes them to bear no influence on the model outcomes discussed in this paper. Nevertheless, details on unstable equilibria of (2.3) can be found in the supplementary material.

### 3.2 Patterned species coexistence

Decreases in resource input cause the spatially uniform states of (2.3) to lose stability to spatio-temporal patterns. Patterns either feature both consumer species or consist of the colonizer species only. Single-species patterns of the locally superior species are always unstable to the introduction of the coloniser species, despite being stable in the absence of any competitor. Significantly, a solution’s species composition is not necessarily conserved across a transition from a spatially uniform to a spatially patterned state. For example, if intraspecific competition of the locally superior species is insufficient to stabilise spatially uniform coexistence, the spatially uniform single-species state nevertheless loses stability to a coexistence pattern (Fig. 3.2 (b)).

Moreover, resource input also determines if a solution represents a patterned ecosystem in which consumers separate into densely populated areas and regions that are left uncolonised (e.g. vegetation patterns) or a spatially non-uniform but nevertheless continuously colonised ecosystem (e.g. savannas). The former state is only attained under severe environmental stress and yields solutions in which consumer densities oscillate between a level of high biomass and zero. By contrast, larger resource input levels result in states in which consumers oscillate between two non-zero biomass levels. Crucially, coexistence in a patterned ecosystem state under low resource input is only possible if intraspecific competition among the coloniser species is sufficiently strong (Fig. 3.2 (a) and (b)). If self-limitation of the coloniser species is weak, then the beneficial effects of its colonisation abilities outweigh the local superiority of its competitor for a wider range of resource input volumes. The destabilisation of coexistence thus occurs at higher resource input levels, where solutions represent spatially non-uniform but continuously colonised states (Fig. 3.2 (c) and (d)). Indeed, if the modelling framework does not account for any intraspecific competition dynamics other than those for the limiting resource (a case that is comparable with that depicted in (Fig. 3.2 (d)), then a spatially non-uniform state in which biomass fluctuates between two nonzero levels is the only possible mode of coexistence (Eigentler and Sherratt 2020c).

Under high-volume resource input regimes, a unique spatially uniform stable state exists for each value of the resource input parameter *A*, but multistability of stable equilibria occurs for lower resource input amounts. Firstly, if a spatial pattern is a stable model outcome for a given precipitation volume, then a range of other patterned solutions are also stable. Such patterns differ in their wavelengths (distance between two consecutive biomass peaks) and their migration speeds (Fig. 3.2 and Fig. 3.3 (a)). Secondly, multistability of single-species and coexistence patterns, as well as coexistence patterns and spatially uniform states also occurs, especially if intraspecific competition among the coloniser species is strong (Fig. 3.2 (a) and (b)).

**Figure 3.3:**
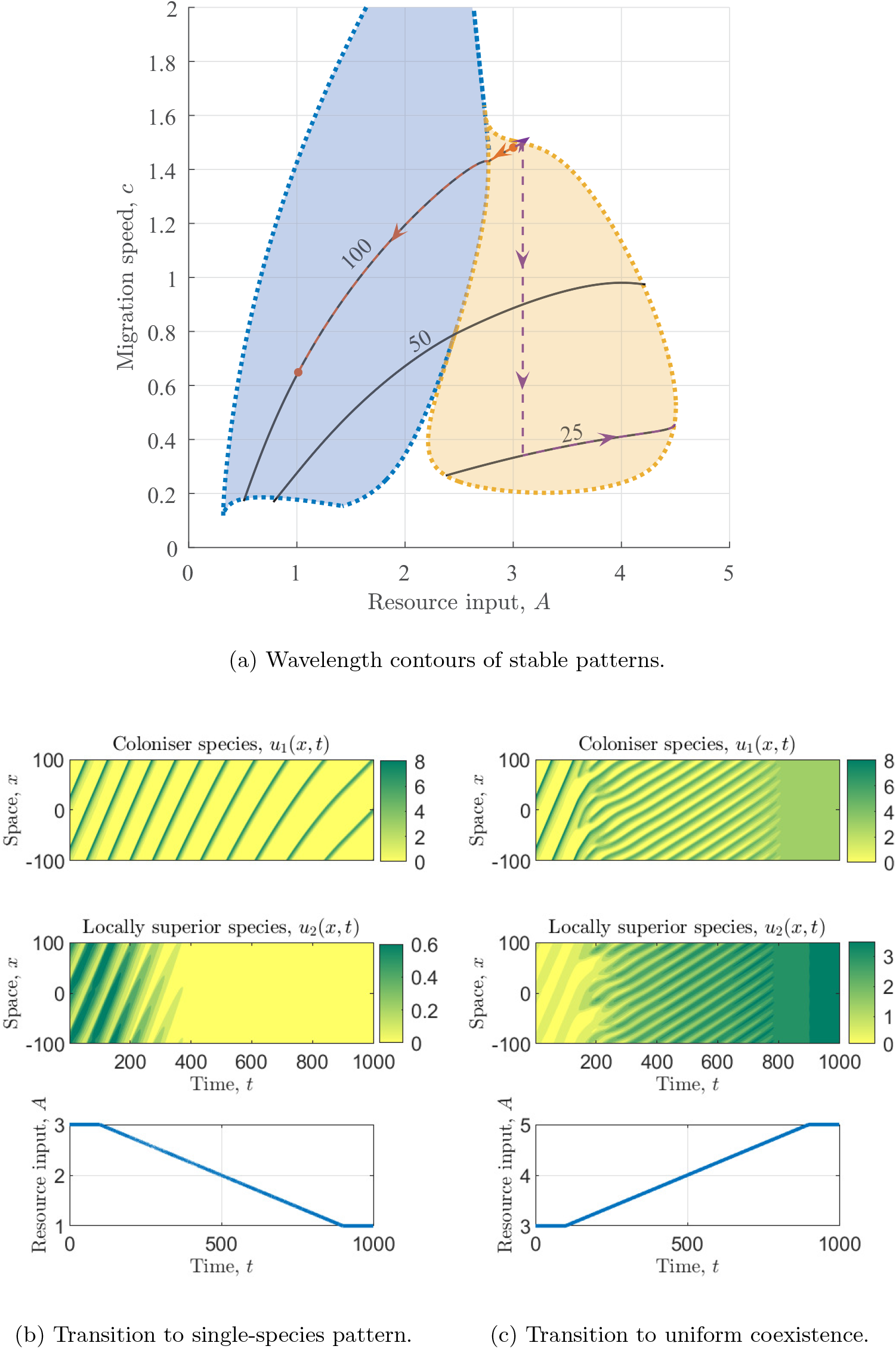
State transitions under changing environmental conditions. Transitions from patterned coexistence to a single-species pattern (b) and a uniform coexistence state (c) under changing resource input volumes are shown. Both model solutions are initiated at resource input volume *A* = 3 and a wavelength of *L* = 100. Solutions follow wavelength contours (solid black curves in (a)) until they are destabilised and a pattern of a new stable wavelength is chosen. Red dashed lines in (a) indicate the dynamics of the solution shown in (b); purple dashed lines that of the model outcome visualised in (c). Note that due to the gradual change in the resource input parameter *A*, patterns lose their periodicity in time. Parameter values correspond to those used in Fig. 3.2 (a).

This singles out patterns’ wavelengths and migration speeds as crucial pieces of information in the understanding of transitions between species coexistence and single-species states. Typically, if a patterned state of a given wavelength loses its stability due to changes in resource input, a transition, such as that depicted in Fig. 3.3 (c), to a pattern of a different wavelength occurs. The multistability of patterned states implies that the newly attained state is not necessarily close to destabilisation. Hence, such transitions cannot typically be undone by simply reversing resource input back to its original level.

A change in wavelength is not the only possible result of a pattern’s destabilisation. If intraspecific competition of the coloniser species is strong, a large range of coexistence patterns undergo a transition to a single-species pattern (of the coloniser) with the same wavelength as environmental stress increases (Fig. 3.2 (a) and (b) and Fig. 3.3). Such a bifurcation is characterised by a stability change of the single-species pattern to the introduction of a competitor species (Eigentler and Sherratt 2020c). Moreover, unlike other bifurcations in the system, such an extinction event can be detected in advance. Decreases in resource input lead to a gradual decrease in the density of the locally superior species until, at the bifurcation, its amplitude approaches zero and the species vanishes from the system. This type of extinction event occurs despite the stability of coexistence patterns at different wavelengths for the same environmental conditions. Thus, species richness in the system not only depends on resource input volume and species properties but also on past states of the ecosystem. In particular, the resilience of coexistence patterns to increases in environmental stress depends on their wavelength and uphill migration speed. The phenomena outlined above are examples of hysteresis, a well-known feature of theoretical models of patterned ecosystems (Sherratt 2013a).

Spatially uniform equilibria and regular spatio-temporal patterns are not the only stable outcomes of the model system (2.3). In the absence of strong intraspecific competition, no direct switch from regular coexistence pattern to a uniform state of the locally superior species occurs (Fig. 3.2 (d)). Instead, this transition takes place via a coexistence state, which lacks any regularity in space and time (example solution is shown in the supplementary material). Currently, I am unable to make any conclusive statements about this type of model solution and its ecological implications, but note that the mappings of stability regions (Fig. 3.2) emphasise that its occurrence relies on the intraspecific competition dynamics of both species being weak. Moreover, numerical simulations suggest that this type of solution only occurs if the local average fitness difference (in the absence of spatial dynamics) *FB*_1_ −*B*_2_ is sufficiently small, despite the coexistence mechanism presented in this paper not relying on such a condition (Eigentler and Sherratt 2020c).

Similar to the spatially uniform states discussed in Sec. 3.1, stable patterned model outcomes of (2.3) are also complemented by unstable solutions. While in this context, unstable states do not have any ecological relevance, they are essential to understand the full solution structure of the mathematical model. Examples of bifurcation diagrams as well as more information on the bifurcations that occur in the system are given in the supplementary material.

### 3.3 Consumer species’ distribution

Depending on system parameters, the multispecies model (2.3) captures the spatial species distribution within single bands that is, for example, being reported from dryland vegetation patterns (Seghieri et al. 1997). In other words, the stripe’s edge intercepting the unidirectional resource flux is dominated by the coloniser species, while the locally superior species is typically found in the centre and towards the down-flux region of each band (Fig. 3.4b).

**Figure 3.4:**
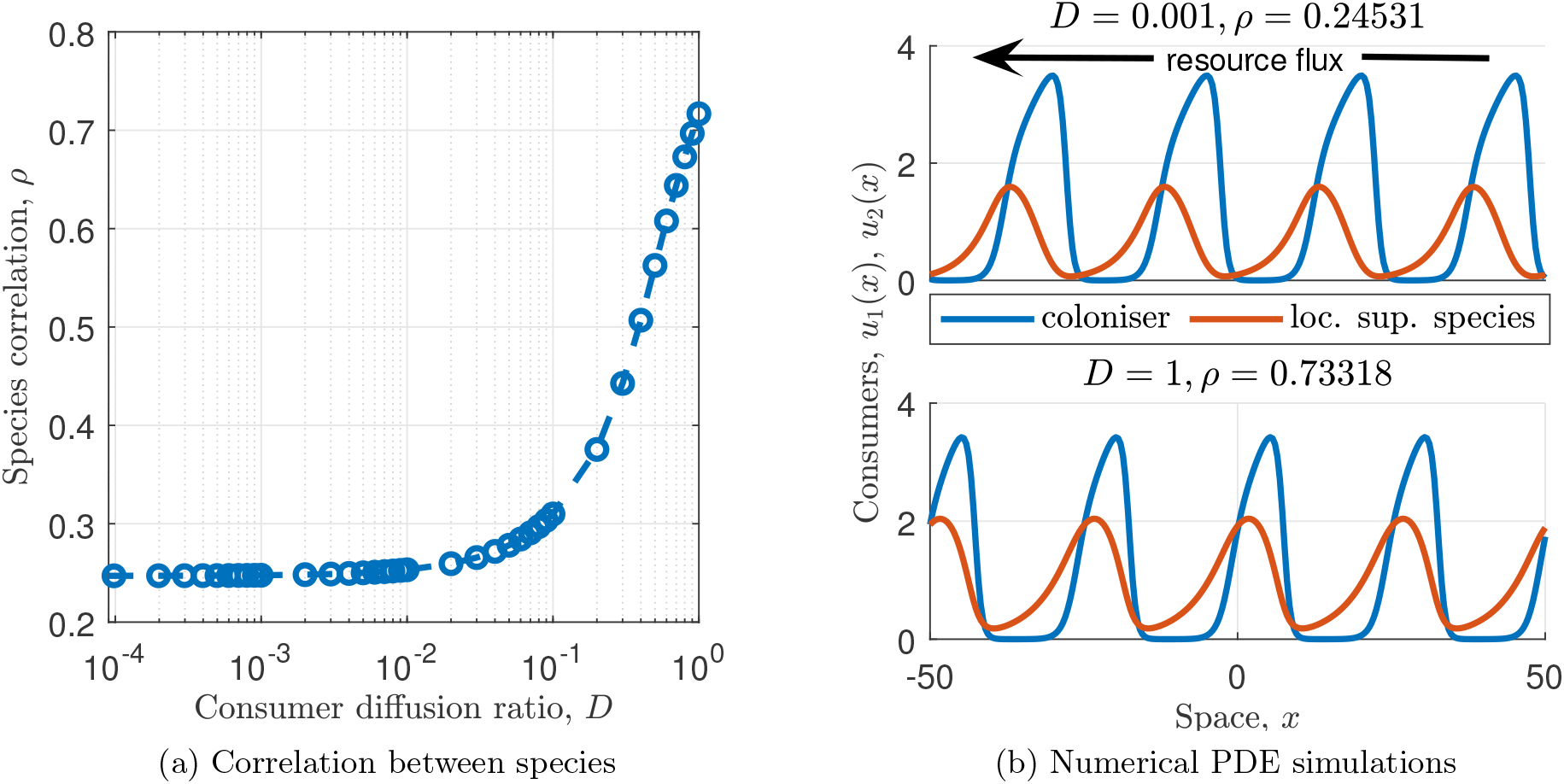
The spatial correlation between the two consumer species components of model solutions under changes in the diffusion coefficient *D* are shown in (a). Part (b) visualises solution profiles (*u*_1_,*u*_2_ only) for specific values of *D*. Correlation increases as dispersal behaviour becomes more similar. Other parameter values are *B*_1_ = 0.45, *B*_2_ = 0.049, *F* = *H* = *D* = 0.11, *ν* = 182.5, *d* = 500, *k*_1_ = *k*_2_ = 5 and *A* = 2.

The ratio of the consumer species’ diffusion coefficients *D* has the most significant impact on the correlation between both plant species (Fig. 3.4). If the species with slower growth rate also disperses at a slower rate, then the correlation between both species is small, as the uphill edge of each vegetation band features a high density of the faster disperser only. As the difference between the species’ diffusion coefficients becomes smaller, the correlation between the consumer densities increases. For *D* = 1, i.e. when both species diffuse at the same rate, both species feature near the top edge of each stripe at a high density. Nevertheless, solution components are not exactly in phase due to the faster growth and mortality dynamics of the coloniser species. (Fig. 3.4b).

## 4 Discussion

The exploration of coexistence mechanisms that prevent competitive exclusion in resource limited ecosystems has been a common research topic in theoretical ecology for many decades (Chesson 2000; McPeek 2012; Tilman 1982; Volterra 1928). The importance of considering intraspecific competition dynamics has been established early in the history of coexistence theory (e.g. (R. MacArthur 1970)). Self-limitation of species enables coexistence because it keeps their abundances sufficiently low to prevent resource monopolisation by any one species (Chesson 2000; Tilman 1982). More recently, heterogeneities caused by spatial self-organisation in otherwise homogeneous environments have been singled out as an alternative potential explanation of species coexistence under competition for a sole limiting resource (Cornacchia et al. 2018; Eigentler and Sherratt 2020c). Interestingly, these processes are typically associated with opposite impacts on population growth in general: intraspecific competition corresponds to negative density-dependence, but spatial self-organisation involves local facilitation.

Previous theoretical studies suggest that spatial self-organisation on its own enables species coexistence under relatively favourable environmental conditions, if there exists a trade-off between species’ colonisation and dispersal abilities (Eigentler and Sherratt 2020c). In contrast to classical coexistence theory, coexistence under such a trade-off can occur without spatial segregation of species (Gravel et al. 2010; Hassell et al. 1994; Horn and R. H. MacArthur 1972; Levins and Culver 1971; Tilman 1994). Nevertheless, pattern formation on its own has been unable to explain species coexistence across a wide range of the resource input gradient (Eigentler and Sherratt 2020c; Pueyo et al. 2010) and has, in particular, failed to capture coexistence in patterned ecosystems where consumer species separate into colonised and uncolonised patches under severe environmental stress. In this paper, I show that the combination of facilitative self-organisation and self-limiting intraspecific competition in a theoretical framework provides more insights into the coexistence of a coloniser species with a locally superior species in states ranging from spatially uniform consumer abundance to regular striped patterns with uncolonised interband regions.

A widely applicable result of coexistence theory is that coexistence of two species can occur if each species’ intraspecific competition is stronger than its impact on the competitor species (e.g. (Chesson 2000; McPeek 2012)). Interestingly, the combination of intraspecific competition with spatial self-organisation dynamics show that different species’ self-limitation is the facilitator of coexistence at opposite ends of the resource input spectrum. This phenomenon arises despite the assumption that both species do not qualitatively differ from each other in any of their functional responses to the environment. Under high resource availability, for which consumer species can attain a spatially uniform state, strong intraspecific competition of the locally superior species facilitates coexistence (Fig. 3.1). If self-limitation of the locally superior species is sufficiently strong, it cannot utilise all of the available resource and thus allows for coexistence with a second species (McPeek 2012). In this context, local superiority corresponds to a higher local average fitness, which, in my theoretical framework, is determined by the stability of spatially uniform single-species equilibria. While in the multispecies model presented in this paper, the stability regions of these steady states do not overlap, complications in the definition of the locally superior species may arise in other modelling frameworks if bistability of single-species states occurs.

The impact of intraspecific competition changes significantly if resource availability is low. Under such environmental conditions, the spatial self-organisation principle destabilises spatially uniform states and leads to pattern formation. Firstly, species coexistence requires the locally superior species to be inferior in its colonisation abilities (Eigentler and Sherratt 2020c). This balance enables species coexistence due to spatial heterogeneities in the environment, caused by the spatial self-organisation. The potential of intercepting the resource flux in uncolonised regions is high, which can be exploited by the coloniser. The redistribution of the limiting resource towards areas of high biomass consequently facilitates the growth of the second species. Eventually, the locally superior species locally outcompetes the superior coloniser, thus creating a balance that facilitates coexistence. Secondly, coexistence in patterned form is facilitated by strong intraspecific competition of the coloniser species (Fig. 3.2 (b)). In contrast to the coexistence mechanism which applies under high resource availability and is discussed above, the intraspecific competition’s impact on resource availability is not the cause of coexistence. Instead, coexistence under severe resource scarcity is enabled by the strengthening of the facilitative balance between the two species’ colonisation abilities and local competitiveness. In the absence of intraspecific competition, the coloniser is able to tip that balance in its favour by being able to colonise new areas faster than being outcompeted locally by its competitor. Strong self-limitation of the coloniser prevents this and thus facilitates coexistence.

My model analysis further highlights the importance of hysteresis in the understanding of species coexistence in self-organised ecosystems. As is common with pattern-forming systems, multistability of equilibria occurs. In other words, given any set of parameters describing consumer species and environmental conditions, no conclusive statement about the model outcome can be made. Instead, information about a pattern’s history is required to obtain more information about its future dynamics. In particular, tipping points or other significant transitions, such as extinction events, cannot be pinned down to a specific level of resource without having information about the ecosystem’s current and past states (Fig. 3.3). Theoretical studies of hysteresis in patterned vegetation in the context of only one species highlight that this property could be of crucial importance to advance our understanding of the ecosystem dynamics in general, not only of the transitions between coexistence and single-species states (Dagbovie and Sherratt 2014; Sherratt 2013a). Hysteresis is believed to be of crucial importance in many ecosystems (United Nations Convention to Combat Desertification 2017), but empirical evidence of hysteresis in patterned ecosystems is challenging to detect (Deblauwe et al. 2011) and only limited empirical data on the phenomenon exists (Trichon et al. 2018).

A novelty of results presented in this paper is the proposal of a coexistence mechanism for resource-limited patterned ecosystems that does not rely on species-specific assumptions. In the context of vegetation patterns, previous theoretical models have only captured species coexistence by imposing qualitative differences on the competing species. For example, Baudena and Rietkerk 2013 and Nathan et al. 2013 highlight that species coexistence can occur if only one species contributes to the system’s pattern-forming feedback loop. In the the modelling framework (2.3), this would correspond to *H* = 0. While this parameter choice indeed leads to stable coexistence in (2.3), my analysis highlights that such a restriction is not necessary for species coexistence to occur. Nevertheless, I speculate that qualitative differences in the short-range facilitation by consumer species (i.e. different nonlinear dependencies on *u*_1_ and *u*_2_ in the facilitation term of the per capita growth rate (2.2) instead of the linear responses in *u*_1_ + *Hu*_2_) may significantly affect the coexistence mechanism, due to its impact on the colonisation abilities of both species. In particular, different nonlinear responses may alter which species is superior in its colonisation abilities depending on environmental conditions and population levels, thus breaking the coexistence mechanism presented in this paper. Similarly, adaptation to different soil moisture niches has been singled out as a potential coexistence mechanism in the context of dryland vegetation patterns (Callegaro and Ursino 2018; Ursino and Callegaro 2016). This could be accounted for by different functional responses of species’ growth rates to the resource density in my modelling framework. These approaches present promising avenues for further exploration of coexistence mechanisms among specific species pairs for which empirical evidence of differing functional responses to the environment is available. By contrast, the coexistence mechanism presented in this paper exclusively relies on quantitative comparative assumptions. As a consequence, it not only applies to a wider combination of plant species but, due to the deliberate simplicity of the theoretical framework, can also be extended to other consumer-resource systems governed by self-organisation principles.

It is worth emphasising that the coexistence mechanism attributed to spatial self-organisation presented in this paper differs from that proposed in a different class of model: rock-paper-scissor-type systems. In those models, coexistence is facilitated by competitive hierarchies of three or more species that follow the idea of the well-known children’s game of the same name. Under such assumptions, spatial self-organisation occurs due to spatial segregation and leads to each coexisting species occupying its own habitat in a larger spatial domain (Avelino et al. 2019; Kerr et al. 2002; Lowery and Ursell 2019; Reichenbach et al. 2007), similar to what is predicted by coexistence theory based on competition-colonisation trade-offs (e.g. (Gravel et al. 2010; Hassell et al. 1994; Horn and R. H. MacArthur 1972; Levins and Culver 1971; Tilman 1994)). By contrast, the theoretical model presented in this paper captures species coexistence within single colonised patches, in agreement with field observations of different patterned ecosystems (Babarro and Comeau 2014; Bekker 2005; Porri et al. 2016; Reise et al. 2017; Seghieri et al. 1997; Valentin et al. 1999). Moreover, it also captures both the migration of consumer bands in the direction opposite to the unidirectional resource flow and the spatial species distribution within single stripes (Fig. 3.4b). Such a movement is reported from both vegetation patterns (Deblauwe et al. 2012; Valentin et al. 1999) and ribbon forests (Bekker 2005) at rates of a few decimetres per year in both cases. I am not aware of any corresponding data for mussel beds as they are typically forced to reassemble regularly after dislodgement due to winter storms. Moreover, the assumption of unidirectional resource flux does not hold in the context of intertidal mussel beds. Unlike on sloped terrain in drylands where water is consistently flowing downhill or at Subalpine tree lines where there is little variation in wind direction causing a consistent shift of snow, algae flow in intertidal regions follows the roughly periodic occurrence of the tide. Indeed, the inclusion of a sinusoidal advection rate in a related modelling framework has shown that only small-scale oscillations of bands occur if a more realistic tidal algae flow is considered (Sherratt and Mackenzie 2016).

Moreover, field observations of banded vegetation patterns report that the top edge of each stripe is dominated by annual grasses (Seghieri et al. 1997), thus coined to be the *coloniser species*, while other species are mostly confined to the centre and lower regions of each stripe. The comprehensive analysis of spatial correlation between both species in coexistence solutions of (2.3) shows that the spatial species distribution in a single consumer band mainly depends on the species’ dispersal behaviour. The faster the diffusion of the coloniser species in relation to its competitor, the more pronounced is its presence at the edge of the stripe that intercepts the unidirectional resource flux. This supports the empirical hypothesis that the pioneering character of grasses in grass-tree coexistence is indeed caused by the faster dispersal of the coloniser species (Seghieri et al. 1997). By contrast, I am not aware of any field data related to this phenomenon from other patterned ecosystems. However, in the context of mussel beds, the characterisation of a species’ colonisation capabilities through its ability to adhere to a substratum, a property that differs among mussel species (Babarro and Comeau 2014), could facilitate future laboratory experiments to test the generality of this hypothesis.

In the model presented in this paper, all intraspecific competition dynamics are combined into one single parameter for each species. To gain a better understanding of how the inclusion of intraspecific competition dynamics in modelling frameworks governed by self-organisation principles affects species coexistence, more information on these dynamics is needed. Promising first steps have been made through the explicit inclusion of (auto-)toxicity effects on interacting plant species (Marasco et al. 2020). A further potential shortfall of the model presented in this paper is that interspecific competition dynamics are restricted to competition for the limiting resource only. In another potential model extension, the self-limitation term in (2.2) could be adjusted to also include additional interspecific competition dynamics. While the impact of the interplay of intraspecific and interspecific competition has been the focus of coexistence theory since its early days (e.g. Volterra 1928), its precise impact on patterned ecosystems remains to be uncovered. The addition of additional interspecific competitive interactions could lead to a separation of competitive dynamics across, for example, the resource input spectrum. In combination with temporal and spatial fluctuations of resources, as induced by spatial self-organisation principles, this could enrich the solution dynamics of the system and thus significantly impact species coexistence (Chesson 2000). These examples suggest potential avenues of further exploration in the context of ecosystems in which both spatial self-organisation and competition dynamics play a significant role.

Finally, I hypothesise that results presented in this paper can be extended to applications in ecosystems that lack the unidirectional resource flux that forms part of the modelling assumptions for (2.3). While this assumption holds true for many different patterned ecosystems, including the examples used throughout the paper, spatial patterns are also known to occur in the absence of unidirectional resource fluxes, such as in “fairy circles” (a common type of vegetation pattern on flat terrain) (Getzin et al. 2014; Getzin et al. 2016). Nevertheless, numerical continuation of model solutions in the advection parameter *ν* to values close to zero, as well as direct numerical integration of the system with *ν* = 0, suggest that my modelling framework also captures consumer coexistence in the absence of unidirectional resource flux. However, I emphasise that a comprehensive treatment of the model in this case would pose a significant challenge, as my model analysis relies on the occurrence of the advection term in (2.3c) (see supplementary material).

Mathematical models can provide powerful tools to advance our understanding of ecosystem functioning and dynamics. In this paper, a model of two consumer species that possess the ability to self-organise into spatial patterns and compete for a limited resource is used to investigate potential modes of coexistence. While the phenomenological and generic nature of the theoretical framework potentially makes it applicable to a wide range of different ecosystems, comprehensive tests to confirm the hypotheses on coexistence mechanisms, species distribution and the role of hysteresis would require comparisons with high-quality field data. It is possible to obtain comprehensive data on properties such as topography (Sugarbaker et al. 2014), movement of stripe patterns (Deblauwe et al. 2012) or historical rainfall data (Sherratt 2015), but other key metrics, such as on species composition or predictions of future environmental conditions are currently unattainable over long temporal and large spatial scales (but at fine resolutions). Nevertheless, the hypotheses put forward by the framework presented in this paper offer a valuable foundation that can both guide future field studies and be tested once advances in technologies such as remote sensing and machine learning make more empirical data on species diversity in stress-induced patterned ecosystems available (Lary et al. 2016).

## Acknowledgements

I thank Jonathan A. Sherratt (Heriot-Watt University) for inspiring discussions and useful comments on the manuscript, Jamie J. R. Bennett (Ben-Gurion University) for helpful discussions and Christopher A. Klausmeier (Michigan State University) for useful comments on earlier versions of the manuscript.

