## Supplementary material for "Species coexistence in resource-limited patterned ecosystems is facilitated by the interplay of spatial self-organisation and intraspecific competition"

Lukas Eigentler[1,2,3]

[1]Division of Molecular Microbiology, School of Life Sciences, University of Dundee, Dundee DD1 5EH, United Kingdom
[2]Division of Mathematics, School of Science and Engineering, University of Dundee, Dundee DD1 4HN, United Kingdom
[3]Maxwell Institute for Mathematical Sciences, Department of Mathematics, Heriot-Watt University, Edinburgh EH14 4AS, United Kingdom

#### 1 Dimensional model and its nondimensionalisation

The nondimensional multispecies model used in the manuscript is obtained from the dimensional model

$$\frac{\partial u_1}{\partial t} = \overbrace{\alpha_1^{(1)} w u_1 \left(\alpha_2^{(1)} u_1 + \alpha_2^{(2)} u_2\right)\left(1 - \frac{u_1}{\alpha_{10}^{(1)}}\right)}^{\text{consumer growth}} - \overbrace{\alpha_3^{(1)} u_1}^{\text{baseline mortality}} + \overbrace{\alpha_5^{(1)} \frac{\partial^2 u_1}{\partial x^2}}^{\text{consumer dispersal}}, \quad (1.1a)$$

$$\frac{\partial u_2}{\partial t} = \overbrace{\alpha_1^{(2)} w u_2 \left(\alpha_2^{(1)} u_1 + \alpha_2^{(2)} u_2\right)\left(1 - \frac{u_2}{\alpha_{10}^{(2)}}\right)}^{\text{consumer growth}} - \overbrace{\alpha_3^{(2)} u_2}^{\text{baseline mortality}} + \overbrace{\alpha_5^{(2)} \frac{\partial^2 u_2}{\partial x^2}}^{\text{consumer dispersal}}, \quad (1.1b)$$

$$\frac{\partial w}{\partial t} = \underbrace{\alpha_6}_{\text{resource input}} - \underbrace{\alpha_7 w}_{\text{natural resource depletion}} \underbrace{- w\left(u_1 + u_2\right)\left(\alpha_2^{(1)} u_1 + \alpha_2^{(2)} u_2\right)}_{\text{resource consumption by consumers}} + \underbrace{\alpha_8 \frac{\partial w}{\partial x}}_{\text{unidirectional resource flux}} + \underbrace{\alpha_9 \frac{\partial^2 w}{\partial x^2}}_{\text{resource diffusion}}. \quad (1.1c)$$

The model is based on the single-species reaction-advection-diffusion model by Klausmeier (Klausmeier 1999) and is an extension of the multispecies models used in (Eigentler and Sherratt 2019; Eigentler and Sherratt 2020). The spatial domain is one-dimensional and $x \in \mathbb{R}$ (units: m) increases in the direction opposite the unidirectional resource flux. Consumer densities at time $t \geq 0$ (years) and $x \in \mathbb{R}$ are denoted by $u_i(x,t)$ (kg m$^{-1}$) and the resource density by $w(x,t)$ (kg m$^{-1}$).

A constant amount of the limiting resource is added to the system per unit time, represented by $\alpha_6$ (kg m$^{-1}$ years$^{-1}$) in the model. Processes independent of the presence of consumers cause a reduction in the resource and are assumed to occur at rate $\alpha_7$ (years$^{-1}$). Resource consumption is the product of three terms: the resource density $w$, the total consumer density $u_1 + u_2$ and a term representing the increase in resource available for consumption in dense biomass patches $\alpha_2^{(1)} u_1 + \alpha_2^{(2)} u_2$. The constants $\alpha_2^{(1)}$ and $\alpha_2^{(2)}$ (both m$^2$ years$^{-1}$ kg$^{-2}$) account for the strength of each species' resource availability enhancement. In the absence of intraspecific competition, growth of each consumer directly corresponds to its resource consumption and $\alpha_1^{(1)}$ and $\alpha_1^{(2)}$ (both dimensionless) quantify the species' resource to biomass conversion ability. Further, consumer growth is limited by density dependent effects, here modelled by a logistic growth-type term with carrying capacities $\alpha_{10}^{(1)}$ and $\alpha_{10}^{(2)}$ (both kg m$^{-1}$), respectively. Consumer mortality occurs at constant rates $\alpha_3^{(1)}$ and $\alpha_3^{(2)}$ ( both years$^{-1}$), respectively. Finally, both consumer densities and the resource density undergo diffusion with coefficients $\alpha_5^{(1)}$, $\alpha_5^{(2)}$ (both m$^2$ years$^{-1}$) and $\alpha_9$ (m$^2$ years$^{-1}$), respectively. If unidirectional resource flux is assumed to occur, then this is modelled by advection with speed $\alpha_8$ (m years$^{-1}$).

A suitable nondimensionalisation of (1.1) is given by

$$u_1 = \left(\frac{\alpha_7}{\alpha_2^{(1)}}\right)^{\frac{1}{2}}\widetilde{u_1}, \quad u_2 = \left(\frac{\alpha_7}{\alpha_2^{(1)}}\right)^{\frac{1}{2}}\widetilde{u_2}, \quad w = \frac{\alpha_7^{\frac{1}{2}}}{\alpha_1^{(1)}\left(\alpha_2^{(1)}\right)^{\frac{1}{2}}}\widetilde{w},$$

$$x = \left(\frac{\alpha_5^{(1)}}{\alpha_7}\right)^{\frac{1}{2}}\widetilde{x}, \quad t = \frac{1}{\alpha_7}\widetilde{t}, \quad A = \frac{\alpha_1^{(1)}\left(\alpha_2^{(1)}\right)^{\frac{1}{2}}\alpha_6}{\alpha_7^{\frac{3}{2}}}, \quad B_i = \frac{\alpha_3^{(i)}}{\alpha_7}, \quad F = \frac{\alpha_1^{(2)}}{\alpha_1^{(1)}},$$

$$H = \frac{\alpha_2^{(2)}}{\alpha_2^{(1)}}, \quad k_i = \frac{\alpha_{10}^{(i)}\left(\alpha_2^{(1)}\right)^{\frac{1}{2}}}{\alpha_7^{\frac{1}{2}}}, \quad D = \frac{\alpha_5^{(2)}}{\alpha_5^{(1)}}, \quad \nu = \frac{\alpha_8}{\left(\alpha_5^{(1)}\alpha_7\right)^{\frac{1}{2}}}, \quad d = \frac{\alpha_9}{\alpha_5^{(1)}}.$$

Substitution into (1.1) and dropping the tildes for brevity gives the nondimensional model studied in the paper, which is

$$\frac{\partial u_1}{\partial t} = \underbrace{u_1\left(w\left(u_1 + Hu_2\right)\left(1 - \frac{u_1}{k_1}\right) - B_1\right)}_{\text{consumer growth/death}} + \underbrace{\frac{\partial^2 u_1}{\partial x^2}}_{\text{consumer dispersal}}, \tag{1.2a}$$

$$\frac{\partial u_2}{\partial t} = \overbrace{u_2\left(Fw\left(u_1 + Hu_2\right)\left(1 - \frac{u_2}{k_2}\right) - B_2\right)} + \overbrace{D\frac{\partial^2 u_2}{\partial x^2}}, \tag{1.2b}$$

$$\frac{\partial w}{\partial t} = \underbrace{A}_{\text{resource input}} - \underbrace{w}_{\text{natural resource depletion}} - \underbrace{w\left(u_1 + u_2\right)\left(u_1 + Hu_2\right)}_{\text{resource consumption by consumers}} + \underbrace{\nu\frac{\partial w}{\partial x}}_{\text{unidirectional resource flux}} + \underbrace{d\frac{\partial^2 w}{\partial x^2}}_{\text{resource diffusion}}. \tag{1.2c}$$

The model parameters are all combinations of different dimensional parameters, but can be interpreted as resource input volume ($A$), consumer mortality rates ($B_1$ and $B_2$), relative growth rate of species $u_2$ compared to species $u_1$ ($F$), consumer-dependent resource availability enhancement due to species $u_2$ ($H$), consumers' maximum standing biomasses per unit area ($k_1$ and $k_2$), relative diffusion coefficient of species $u_2$ compared to that of species $u_1$ ($D$), speed of the unidirectional resource flux ($\nu$) and relative diffusion coefficient of the resource compared to that of species $u_1$ ($d$).

### 2 Methods for model analysis

#### 2.1 Transformation into travelling wave coordinates

Typically, model solutions of (1.2) are either spatially uniform or spatially patterned. In the latter case, they are periodic in space and move in the direction opposite to the unidirectional resource flux of the spatial domain at a constant speed. Thus, spatial patterns in (1.2) belong to the class of periodic travelling waves, an important class of solutions for many reaction-advection-diffusion equations. Periodic travelling waves allow for a coordinate transformation to a comoving frame, as they can be represented by a single variable $z = x - ct$, where $c \in \mathbb{R}$ is the migration speed of the solution profile. This coordinate transformation and setting $u_1(x,t) = U_1(z)$, $u_2(x,t) = U_2(z)$ and $w(x,t) = W(z)$ reduces the PDE system (1.2) to the ODE system

$$WU_1(U_1 + HU_2)\left(1 - \frac{U_1}{k_1}\right) - B_1U_1 + c\frac{\mathrm{d}U_1}{\mathrm{d}z} + \frac{\mathrm{d}^2U_1}{\mathrm{d}z^2} = 0, \tag{2.1a}$$

$$FWU_2(U_1 + HU_2)\left(1 - \frac{U_2}{k_2}\right) - B_2U_2 + c\frac{\mathrm{d}U_2}{\mathrm{d}z} + D\frac{\mathrm{d}^2U_2}{\mathrm{d}z^2} = 0, \tag{2.1b}$$

$$A - W - W(U_1 + U_2)(U_1 + HU_2) + (c + \nu)\frac{\mathrm{d}W}{\mathrm{d}z} + d\frac{\mathrm{d}^2W}{\mathrm{d}z^2} = 0. \tag{2.1c}$$

Spatially patterned solutions of the PDE system (1.2) correspond to limit cycles in the travelling wave ODE system (2.1). If a patterned solution exists in the PDE system (1.2) for a given value of the bifurcation parameter, here the resource input volume $A$, then limit cycles in the ODE system (2.1) exist for a range of different wavespeeds $c$ at different wavelenghts.

#### 2.2 Bifurcation analysis

Properties of spatially uniform single-species states in the model are analytically tractable and an overview is provided below (Sec. 3) and can also be found in (Eigentler 2020). Other properties of single-species states (pattern onset, pattern existence and pattern stability) as well as those of coexistence states (spatially uniform equilibria, pattern onset, pattern existence) are determined through numerical continuation using AUTO-07p (Doedel et al. 2012). Besides standard numerical continuation techniques, this also requires an understanding of the patterns' stability, which can be obtained from the patterns' essential spectra. Those are calculated using a numerical continuation method by Rademacher et al. (Rademacher et al. 2007). I refer to (Rademacher et al. 2007; Sherratt 2012; Sherratt 2013) for full details on the method, and to (Eigentler and Sherratt 2020) for details on the method's implementation for a model similar to (1.2). It is noteworthy that the stability analysis relies on the solutions occurring as travelling waves. In the absence of a unidirectional resource flux (advection), patterned solutions do not move in the direction opposite to the resource flux. In this case, they do not emanate at a Hopf bifurcation on the spatially uniform equilibrium. As the construction of patterned solution based on a continuation from a Hopf bifurcation is a critical part of the numerical continuation algorithm, the method is currently not applicable in the context of no unidirectional resource flux.

### 3 Stability of spatially uniform single-species states

The trivial and semi-trivial biologically relevant spatially uniform equilibria of (1.2) are the extinction steady state $(\overline{u_1^D}, \overline{u_2^D}, \overline{w^D}) = (0,0,A)$, the single-species state

$$\left(\overline{u_1^\pm}, 0, \overline{w_1^\pm}\right) = \left(\frac{A \pm \sqrt{A^2 - 4B_1\left(B_1 + \frac{A}{k_1}\right)}}{2\left(B_1 + \frac{A}{k_1}\right)}, 0, \frac{A}{1 + \left(\overline{u_1^\pm}\right)^2}\right),$$

of the coloniser species $u_1$, which exist provided

$$A > A_{\min}^{(1)} := 2B_1 \left( \frac{1}{k_1} + \sqrt{1 + \frac{1}{k_1^2}} \right),$$

and the single-species state

$$\left(0, \overline{u_2^{\pm}}, \overline{w_2^{\pm}}\right) = \left( 0, \frac{FHA \pm \sqrt{(FHA)^2 - 4B_2H\left(B_2 + \frac{FHA}{k_2}\right)}}{2H\left(B_2 + \frac{FHA}{k_2}\right)}, \frac{A}{1 + H\left(\overline{u_2^{\pm}}\right)^2} \right),$$

of the locally superior species, which exist provided

$$A > A_{\min}^{(2)} := \frac{2B_2}{FH} \left( \frac{1}{k_2} + \sqrt{H + \frac{1}{k_2^2}} \right).$$

Further, a pair of spatially uniform coexistence equilibria $(\overline{u_1^{C,\pm}}, \overline{u_2^{C,\pm}}, \overline{w^{C,\pm}})$ exists. Even though a closed-form expression can be obtained, the equilibria's algebraic complexity makes any analytical approach infeasible and the their properties are instead addressed through numerical continuation techniques.

The stability of trivial and semi-trivial spatially uniform equilibria to spatially uniform perturbations is determined by a standard linear stability approach. The extinction steady state $(\overline{u_1^D}, \overline{u_2^D}, \overline{w^D})$ is always linearly stable (eigenvalues of Jacobian are $-B_1$, $-B_2$, $-1$). The equilibrium $(\overline{u_1}^+, 0, \overline{w_1}^+)$ of the coloniser species is stable for

$$A < A_u^{(1)} := \frac{B_2^2 + k_1^2 (B_2 - FB_1)^2}{Fk_1 (B_2 - FB_1)},$$

provided $0 < B_2 - FB_1 < FB_1$ and $k_1 > \sqrt{B_2(2FB_1 - B_2)}(B_2 - FB_1)^{-1}$, and unstable otherwise. The single-species equilibrium $(\overline{u_1}^-, 0, \overline{w_1}^-)$ is always unstable.

The equilibrium $(0, \overline{u_2^+}, \overline{w_2^+})$ of the locally superior species is stable for

$$A < A_u^{(2)} := \frac{F^2 B_1^2 + Hk_2^2 (B_2 - FB_1)^2}{FHk_2 (FB_1 - B_2)},$$

provided $-B_2 < B_2 - FB_1 < 0$ and $k_2 > \sqrt{B_1 FH(2B_2 - FB_1)}(H(FB_1 - B_2))^{-1}$, and unstable otherwise. The single-species equilibrium $(0, \overline{u_2^-}, \overline{w_2^-})$ is always unstable.

Note that the stability regions of the single-species spatially uniform equilibria never overlap, because $A_u^{(1)} A_u^{(2)} < 0$. The sign of $A_u^i$ is determined by $B_2 - FB_1$, the local average fitness difference between both species in the absence of intraspecific competition (Eigentler and Sherratt 2019). Thus, only the single-species state of the species of higher local average fitness can be stable.

The instabilities that occur as $A$ is increased are due to the introduction of the competitor species, which causes the single-species state to lose stability to the coexistence equilibrium. In the corresponding single-species models, both $(\overline{u_1^+}, 0, \overline{w_1^+})$ and $(0, \overline{u_2^+}, \overline{w_2^+})$ are stable for any resource input value in their existence ranges if $B_i < 2$.

Existence and stability of the coexistence equilibrium $(\overline{u_1^{C+}}, \overline{u_2^{C+}}, \overline{w^{C+}})$ are found using the numerical continuation software AUTO-07p (Doedel et al. 2012) and are presented in the main text. Its complement $(\overline{u_1^{C-}}, \overline{u_2^{C-}}, \overline{w^{C-}})$ is always unstable.

The results presented above allow for a classification of the $k_2$-$A$ parameter plane (i.e. the parameter plane spanned by the parameters describing the strength of intraspecific and interspecific competition, respectively) into stability regions (Fig. 3.1 of the main text) and, in

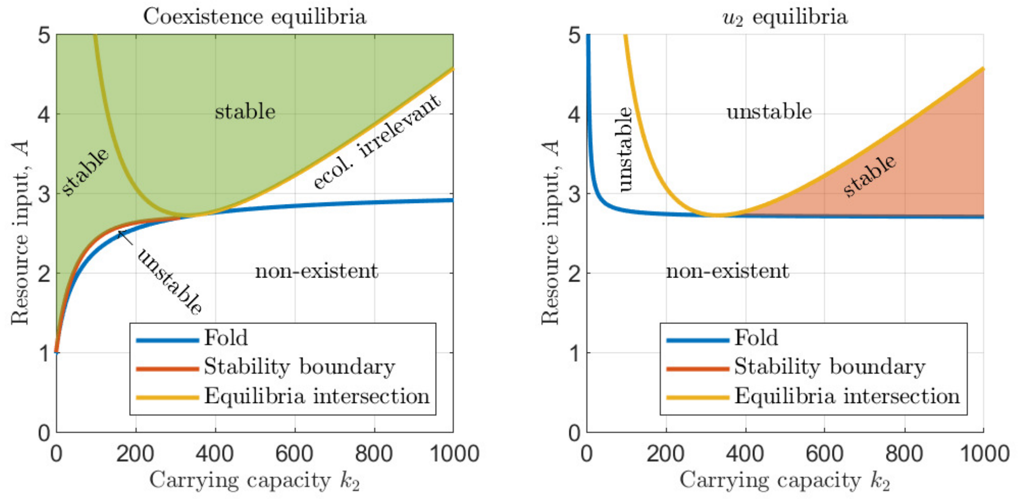


Figure S3.1: Stability and existence regions of equilibria. The left panel shows the parameter regions in which the coexistence equilibrium $(\overline{u_1^{C+}}, \overline{u_2^{C+}}, \overline{w^{C+}})$ is stable (shaded), unstable, ecologically irrelevant (negative densities) and non existent. Similarly, the right panel shows stability, instability and non-existence regions of the single species equilibrium $(0, \overline{u_2^+}, \overline{w_2^+})$ of the superior local competitor $u_2$. Note that the intersection of both equilibria switches from $(0, \overline{u_2^+}, \overline{w_2^+})$ to $(0, \overline{u_2^-}, \overline{w_2^-})$ (and similarly for the coexistence equilibrium) as it passes through the fold of the equilibria. In the right panel, the stability boundary coincides with the equilibria intersection curve to the right of the fold.

more detail, stability and existence regions (Fig. S3.1). Combined, this highlights that the coexistence equilibrium becomes ecologically irrelevant (but continues to exist) as it coincides with the single-species equilibrium, but can also be unstable in the parameter region in which it is positive. Note that the single-species equilibrium becomes unstable at the loci where it intersects with the coexistence equilibrium.

#### 4 Patterned model solutions

The stability diagrams presented in Fig. 3.2 of the main text only provide information on stable model solutions under changes to resource input and intraspecific competition strength. Nevertheless, unstable patterns provide useful information on how different stable states are connected in the system. Thus, a full bifurcation analysis can yield more insights into the model system. Fig. S4.2 in the supplementary material visualises corresponding bifurcation diagrams (for one fixed migration speed only) obtained through numerical continuation. Moreover, the stability diagrams in Fig. 3.2 provide stability information on patterned solutions in the parameter plane spanned by the main bifurcation parameter of the PDE system (resource input, $A$) and the migration speed $c$, which is an emergent property of the PDE system that is made explicit by the transformation into the travelling wave framework (2.1). Hence, it captures the infinite number of different patterned solutions at different migration speeds $c$ that occur due to the infinite length of the spaital domain, but is thus unable to provide any other information on the model solution. To instead visualise the solutions' $L_2$ norm, I fix the migration speed $c$ in the bifurcation diagrams Fig. S4.2. Note that the coexistence equilibrium may exist at negative densities (but never as oscillations about zero that change sign across one period) of the inferior local competitor and this is visualised in Fig. S4.2 by multiplying its $L_2$ norm by $-1$. Further, single species states of $u_1$ are shown as horizontal lines at $u_2 = 0$ in the bifurcation diagrams for $u_2$ and vice versa.

Bifurcation analysis of (1.2) establishes two different mechanisms as the cause of coexistence pattern onset; a Hopf bifurcation of the spatially uniform coexistence equilibrium, and a stability change of a single-species pattern due to the introduction of a competitor species. As a consequence, a coexistence pattern connects a single-species pattern to either the spatially uniform coexistence equilibrium (Fig. S4.2 (a) and S4.2 (c)) or the single-species pattern of its competitor (Fig. S4.2 (b)) in the parameter space.

The onset of spatial patterns via a Hopf bifurcation of a spatially uniform equilibrium is a classical result of bifurcation theory (e.g. (Murray 1989)). Coexistence pattern onset at such a bifurcation is inhibited (i.e. shifted to lower resource input volumes) by both weak intraspecific competition of the superior coloniser and strong intraspecific competition of the locally superior species.

To understand the onset of coexistence patterns from a single-species pattern, information on the latter's stability is required. The stability properties can be split into two distinct and independent mechanisms. To be stable, a single-species pattern requires to be both stable in the absence of a second species and stable to the introduction of its competitor. Onset of coexistence patterns occurs if a single-species pattern loses/gains stability to the introduction of the second species (Eigentler and Sherratt 2020).

A transition between the two mechanisms occurs if intraspecific competition of the locally superior species decreases. Such a decrease reduces the biomass of the locally inferior species in the coexistence equilibrium, and leads to an intersection of the coexistence equilibrium with the single-species equilibrium of the locally superior species. In particular, the Hopf bifurcations on both equilibria coincide for a critical level of the locally superior species' intraspecific competition. This intersection causes a switch in the onset mechanism of coexistence patterns, which hence connect both single-species solution branches for sufficiently weak intraspecific competition of the locally superior species (Fig. S4.2b).

### 5 Other model solutions

Regular spatio-temporal patterns and spatially uniform states are not the only possible model outcomes of the multispecies model (1.2). In the absence of intraspecific competition, there exists an interval of the resource input parameter in which neither of these solution types is stable. Instead, irregular coexistence solutions, such as that shown in Fig. S5.3 are outcomes of the model. My simulations suggest that this type of solution only occurs in a very small region of parameter space.

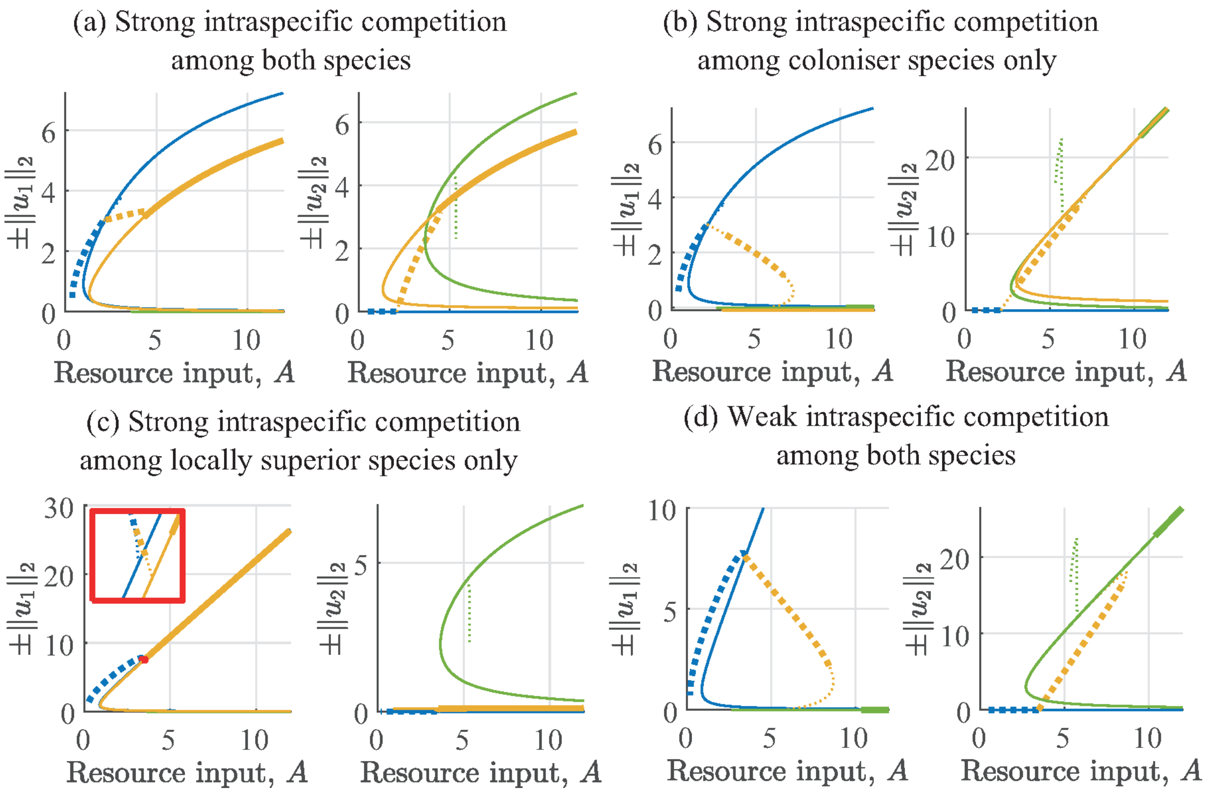


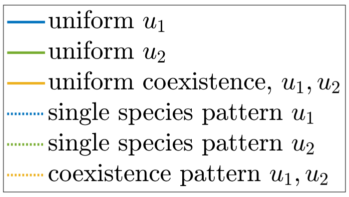


Figure S4.2: Bifurcation diagrams under changing intraspecific competition. Bifurcation diagrams for different values of the carrying capacities $k_1$ and $k_2$ are shown. Solid lines represent spatially uniform states, dotted lines spatial patterns. Bold lines indicate stable states, thin lines correspond to unstable states. Due to the infinite length of the spatial domain, an infinite number of different patterns with different migration speed exist, but only those for one fixed migration speed ($c = 0.42$) are shown. Coexistence states may attain negative densities and thus their $L_2$ norms are multiplied by the density's sign to visualise this in (b). The inset in (c) (axes limits: $A \in [3.35, 3.55]$, $\pm\|u_1\|_2 \in [7.45, 7.65]$) shows a blow-up of the small parameter region in which coexistence pattern occur. The intraspecific competition strengths are $k_1 = k_2 = 10$ in (a), $k_1 = 10, k_2 = 10^4$ in (b), $k_1 = 10^4, k_2 = 10$ in (c) and $k_1 = k_2 = 10^4$ in (d), while other parameter values are $B_1 = 0.45$, $B_2 = 0.05$, $F = H = D = 0.11$, $\nu = 182.5$ and $d = 500$ in (a)-(d).

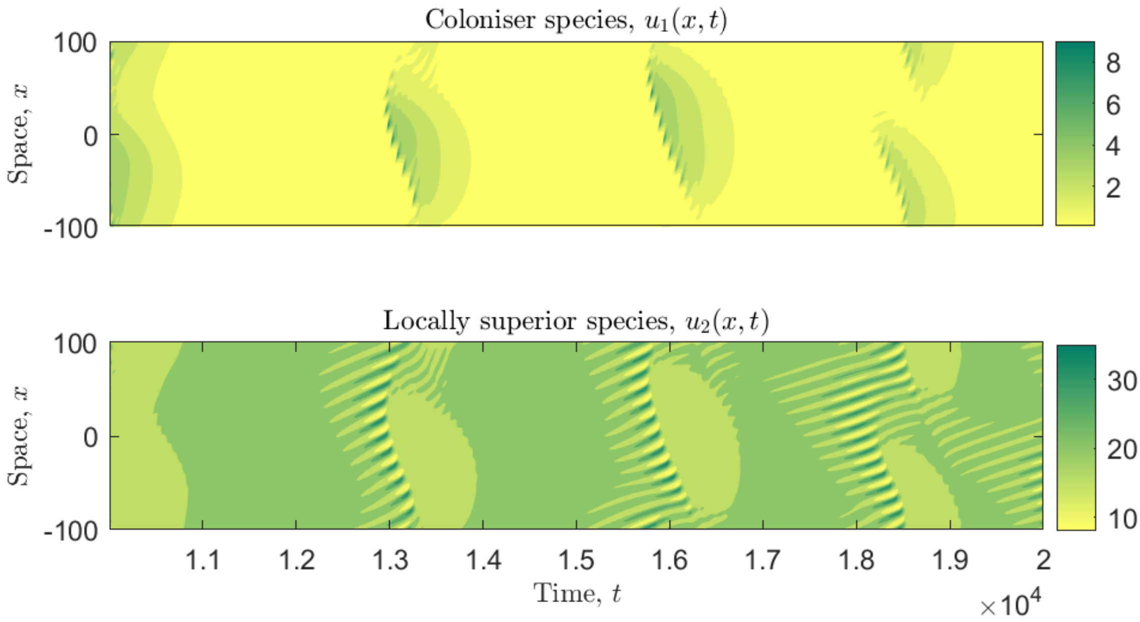


Figure S5.3: Transition between regular coexistence patterns and spatially uniform solution under weak intraspecific competition among both species. This figure shows the long-term behaviour of a model run for the parameter set $A = 9.5$, $B_1 = 0.45$, $B_2 = 0.05$, $k_1 = k_2 = 10^4$, $F = H = 0.11$, $\nu = 182.5$ and $d = 500$. This corresponds to the parameter region shown in Fig. 3.2 (d) of the main text in which neither a regular spatio-temporal nor a spatially uniform equilibrium is stable.
